# Inhibition of PIKfyve kinase prevents infection by Zaire ebolavirus and SARS-CoV-2

**DOI:** 10.1101/2020.04.21.053058

**Authors:** Yuan-Lin Kang, Yi-Ying Chou, Paul W. Rothlauf, Zhuoming Liu, Timothy K. Soh, David Cureton, James Brett Case, Rita E. Chen, Michael S. Diamond, Sean P. J. Whelan, Tom Kirchhausen

## Abstract

Virus entry is a multistep process. It initiates when the virus attaches to the host cell and ends when the viral contents reach the cytosol. Genetically unrelated viruses can subvert analogous subcellular mechanisms and use similar trafficking pathways for successful entry. Antiviral strategies targeting early steps of infection are therefore appealing, particularly when the probability for successful interference through a common step is highest. We describe here potent inhibitory effects on content release and infection by chimeric VSV containing the envelope proteins of Zaire ebolavirus (VSV-ZEBOV) or SARS-CoV-2 (VSV-SARS-CoV-2) elicited by Apilimod and Vacuolin-1, small molecule inhibitors of the main endosomal Phosphatidylinositol-3-Phosphate/Phosphatidylinositol 5-Kinase, PIKfyve. We also describe potent inhibition of SARS-CoV-2 strain 2019-nCoV/USA-WA1/2020 by Apilimod. These results define new tools for studying the intracellular trafficking of pathogens elicited by inhibition of PIKfyve kinase and suggest the potential for targeting this kinase in developing small-molecule antivirals against SARS-CoV-2.

## INTRODUCTION

Membrane-enveloped viruses deliver their contents to cells via envelope protein-catalyzed membrane fusion. Binding of virus to specific host cell receptor(s) triggers membrane fusion, which can occur directly at the plasma membrane or following endocytic uptake. Viruses that require endocytic uptake can use different initial trafficking routes to reach the site of membrane fusion. In endosomes, acidic pH serves to triggers conformational rearrangements in the viral envelope proteins that catalyze membrane fusion, as seen for influenza A virus (IAV) and vesicular stomatitis virus (VSV). For Zaire ebolavirus (ZEBOV), proteolytic processing of the envelope protein by host cell proteases (1) is necessary to expose the receptor binding domain prior to engagement of Niemman-Pick disease type 1C (NPC1 or NPC Intracellular Cholesterol Transporter 1) – the late endosomal-lysosomal receptor protein (2). Proteolytic processing is also required for severe acute respiratory syndrome coronavirus (SARS-CoV) (3, 4), and for the current pandemic SARS-CoV-2 (5). Lassa fever virus (LASV) uses a different mechanism, binding alpha-dystroglycan at the plasma membrane (6), for internalization with a subsequent pH-regulated switch that leads to engagement of lysosomal associated membrane protein 1 (LAMP1) for membrane fusion (7). Lymphocytic choriomeningitis virus (LCMV) also uses alpha-dystroglycan (6) and is internalized in a manner that depends on endosomal sorting complexes required for transport (ESCRT) proteins (8), although it remains unknown whether a second receptor is required.

A hallmark of the endolysosomal system is controlled dynamic trafficking of vesicular carriers among its various sub-compartments. Phophoinositides are markers for defining the identity of these sub-compartments because they are restricted in their distribution to specific intracellular membranes [reviewed in (9)]. Although it is one of the least abundant of the phosphoinositides in cells, PI(3,5)P2 is particularly important for endomembrane homeostasis. It is produced by PIKfyve, which phosphorylates the D-5 position in phosphatidylinositol-3-phosphate (PI3P) to yield phosphatidylinositol 3,5-bisphosphate (PI(3,5)P2) (10). First cloned as mammalian p235 (11), PIKfyve is a 240 kDa class III lipid kinase, present on the cytosolic face of endosomal membranes (12, 13) as part of a ternary complex with the PI(3,5)P2 5-phosphatase Sac3 and ArPIKfyve (14).

Ablation of PIKfyve function by genetic (12, 15) or pharmacological means (16-20) causes endosomal swelling and vacuolation of late endosomes and endolysosomes. It is thought that these changes result from decreased membrane fission and concomitant interference in endosomal traffic (13, 21). Small-molecule inhibitors of PIKfyve, all of which have some structural resemblance to each other, have been studied as potential drugs for treating cancer and autoimmune diseases. These inhibitors include Apilimod (19), Vacuolin-1 (18), a series of 30 Vacuolin-related molecules (22), YM201636 (16), and WX8 chemical family members (20). Physiological effects of these compound in cells include inhibition of autophagy (17, 22, 23), reduced generation of IL-12/IL-23 (24), and reduced dendritic cell infiltration in psoriasis (25).

Apilimod also inhibits infection by several viruses, including ZEBOV. Although it does not alter the pH of endosomes nor inhibit cathepsin B or L (26), Apilimod blocks entry of ZEBOV and other pathogenic filoviruses (27). Several groups reported that Apilimod prevents colocalization of VSV-ZEBOV pseudoviruses with the ZEBOV endosomal receptor NPC1, but does not prevent colocalization with early endosomal antigen 1 (EEA1) (5, 27, 28). Apilimod also inhibits entry of pseudotyped viruses bearing the spike proteins of MERS-CoV, SARS-CoV, and SARS-CoV-2, as well as of authentic mouse hepatitis virus (MHV) particles (5).

Here, we have studied the effects of Apilimod on infection of VSV-eGFP-SARS-CoV-2 and VSV-eGFP-ZEBOV chimeras and showed that Apilimod blocks infection of both, with an IC50 of ∼50 nM. Apilimod and Vacuolin-1 also prevented entry and infection of VSV-MeGFP-ZEBOV and many of the internalized VSV-MeGFP-ZEBOV virions colocalized with NPC1 in the distended, vacuolated endosomes. This suggests that blocking PIKfyve kinase has the same downstream effects on these viruses, even though VSV-eGFP-SARS-CoV-2 does not require interaction with NPC1 for membrane fusion. Apilimod also inhibits infection by authentic SARS-CoV-2 strain 2019-nCoV/USA-WA1/2020 virus, with an IC50 slightly lower than its IC50 for the VSV-eGFP-SARS-CoV-2. We suggest that Apilimod, which has passed safety tests in previous human clinical trials for non-viral indications (24, 25, 29, 30), is a potential starting point for developing small-molecule entry inhibitors of SARS-CoV-2 that could limit infection and disease pathogenesis.

## RESULTS

### Apilimod inhibits infection of VSV-MeGFP-LCMV and VSV-ZEBOV

We inoculated SVG-A cells with vesicular stomatitis virus (VSV) chimeras expressing the viral matrix protein (M) fused to eGFP (MeGFP). The chimeras include VSV (VSV-MeGFP, which initiates fusion at pH<6.2), VSV-V269H GP (VSV-MeGFP-V269H, a variant of VSV GP that initiates fusion at pH<5.8), rabies virus GP (VSV-MeGFP-RABV), Lassa virus GP (VSV-MeGFP-LASV), lymphocytic choriomeningitis virus GP (VSV-MeGFP-LCMV) or Zaire Ebola virus GP (VSV-MeGFP-ZEBOV). Following the incubation protocol summarized in **Fig 1A**, we tested the effects on infection of Apilimod or Vacuolin-1; both compounds are small-molecule inhibitors of PIKfyve kinase, which generates PI(5)P and PI(3,5)P2 in late endosomes and lysosomes. Using a flow cytometry based-assay to monitor a single round of infection determined by expression of viral MeGFP (**Fig. 1B**), we found that Apilimod and Vacuolin-1 potently inhibit VSV-MeGFP-ZEBOV infection (**Fig. 1C**). These results agree with results obtained by others with Apilimod (26, 31) in different cell types infected with MLV virus pseudotyped with ZEBOV GP or with Ebola virus itself (26, 27, 32). Apilimod was a less effective inhibitor of VSV-MeGFP-LCMV infection, and Vacuolin-1 had no effect at the concentration used. In contrast, Apilimod and Vacuolin-1 failed to prevent infection by VSV-MeGFP, VSV-MeGFP-V269H, VSV-MeGFP-RABV, or VSV-MeGFP-LASV (**Fig. 1C**). IN1 (33), an inhibitor of the phosphoinositide kinase Vps34, the main endosomal generator of PI3P, also interfered with VSV-MeGFP-LCMV and VSV-MeGFP-ZEBOV infection (**Fig. 1C**). All of these viruses require low pH to trigger viral membrane fusion with the endosomal membranes, and as expected, infection was fully blocked by Bafilomycin A1, which inhibits the vacuolar type H^+^-ATPase (V-ATPase) acidification activity (**Fig. 1C**).

**Figure 1.**
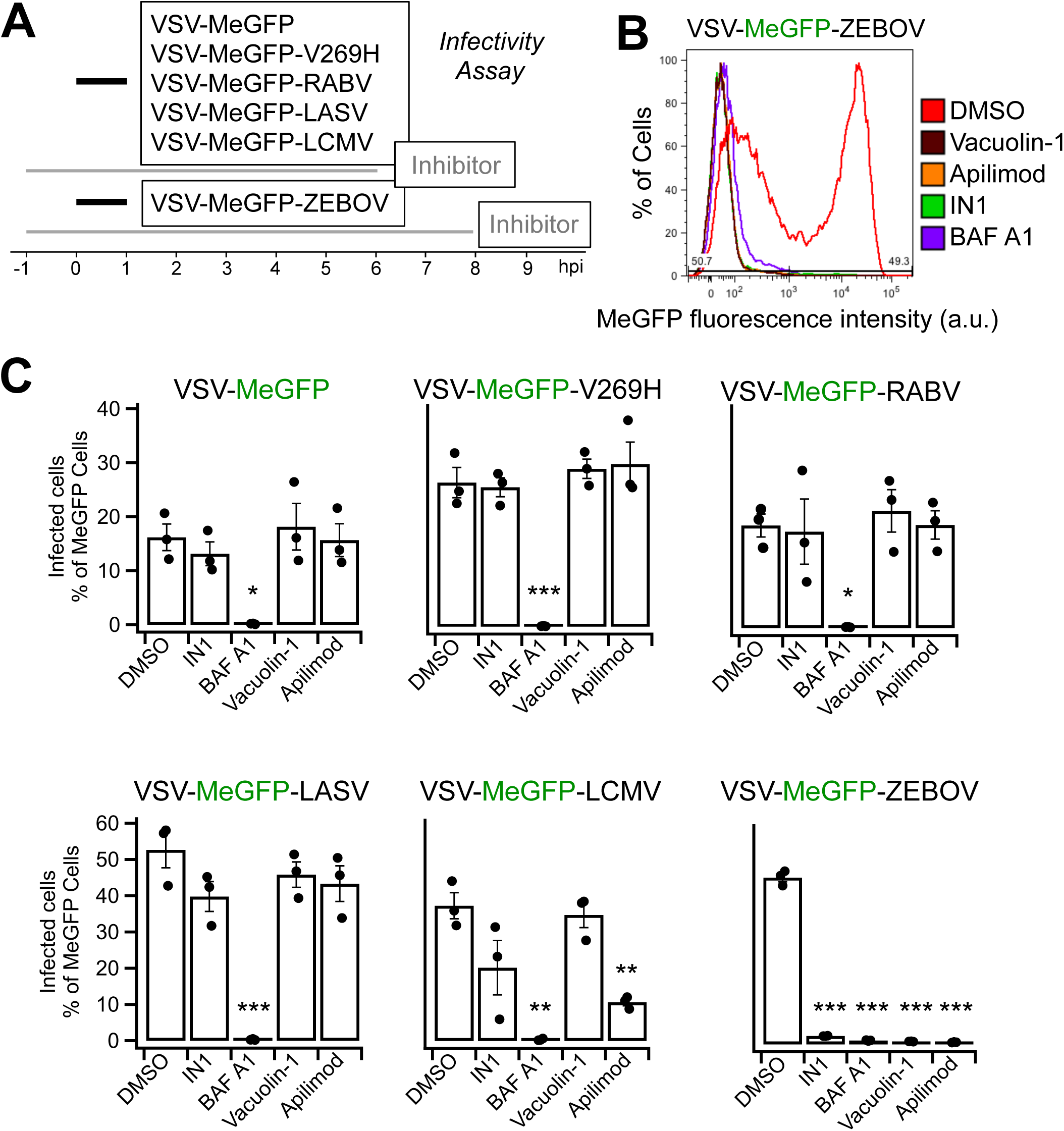
Apilimod and Vacuolin-1 inhibit VSV-MeGFP-ZEBOV infection. **(A)** Schematic of infectivity assay, where SVG-A cells were pretreated for 1 h with 5 µM Vacuolin, 5 µM Apilimod, 5 µM IN1, or 10 nM BAF A1 and subsequently infected with VSV-MeGFP (multiplicity of infection, MOI = 2), VSV-MeGFP-V269H (MOI = 1), VSV-MeGFP-RABV (MO I= 0.6), VSV-MeGFP-LASV (MOI = 0.6), VSV-MeGFP-LCMV (MOI = 0.6) or VSV-MeGFP-ZEBOV (MOI = 0.6) for 1 h in the presence of drugs. The cells were then washed to remove unbound virus and incubated for the indicated times in the presence of drugs. The cells were then fixed and the percentage of cells expressing viral MeGFP was measured by flow cytometry. **(B)** Representative flow cytometry results of an infection assay using VSV-MeGFP-ZEBOV. **(C)** Quantification of the infectivity is shown with averages from three independent experiments per condition each determined as a duplicate measurement (error bars show SEM). The statistical significance was determined using a one-way ANOVA and Tukey *post-hoc* test (*, P ≤ 0.05; **, P ≤ 0.01; ***, P ≤ 0.001).

### Apilimod and Vacuolin-1 prevent cytoplasmic entry of VSV-MeGFP-ZEBOV

Productive infection requires delivery of the viral ribonucleoprotein core (RNP) into the cytosol. In these experiments, we deemed RNP delivery, as monitored by single cell fluorescence microscopy imaging (experimental protocol summarized in **Fig. 2A** and **3A**), to be successful when fluorescent MeGFP encapsulated in the incoming virus appeared at the nuclear margin of infected cells. The representative examples of VSV infection and RNP core release shown in **Fig. 2B** were obtained in the absence or presence of cycloheximide, which prevents viral protein expression. In the absence of cycloheximide (*left panel*), large amounts of newly synthesized MeGFP are present throughout the cell. In the presence of cycloheximide (*right panel*), we observed MeGFP in virions (fluorescent spots) as well as released MeGFP concentrated at the nuclear margin. We scored the effect of Apilimod, Vacuolin-1 or IN1 on RNP delivery by VSV-MeGFP, VSV-MeGFP-V269H and VSV-MeGFP-ZEBOV by determining the appearance of MeGFP at the nuclear margin in cycloheximide-treated cells. Consistent with the infection results, Apilimod, Vacuolin-1 and IN1 prevented entry of VSV-MeGFP-ZEBOV but not of VSV-MeGFP or VSV-MeGFP-V269H. As expected, Bafilomycin A1 blocked entry of all viruses (images in **Fig. 2C** and quantification in **Fig. 2D**).

**Figure 2.**
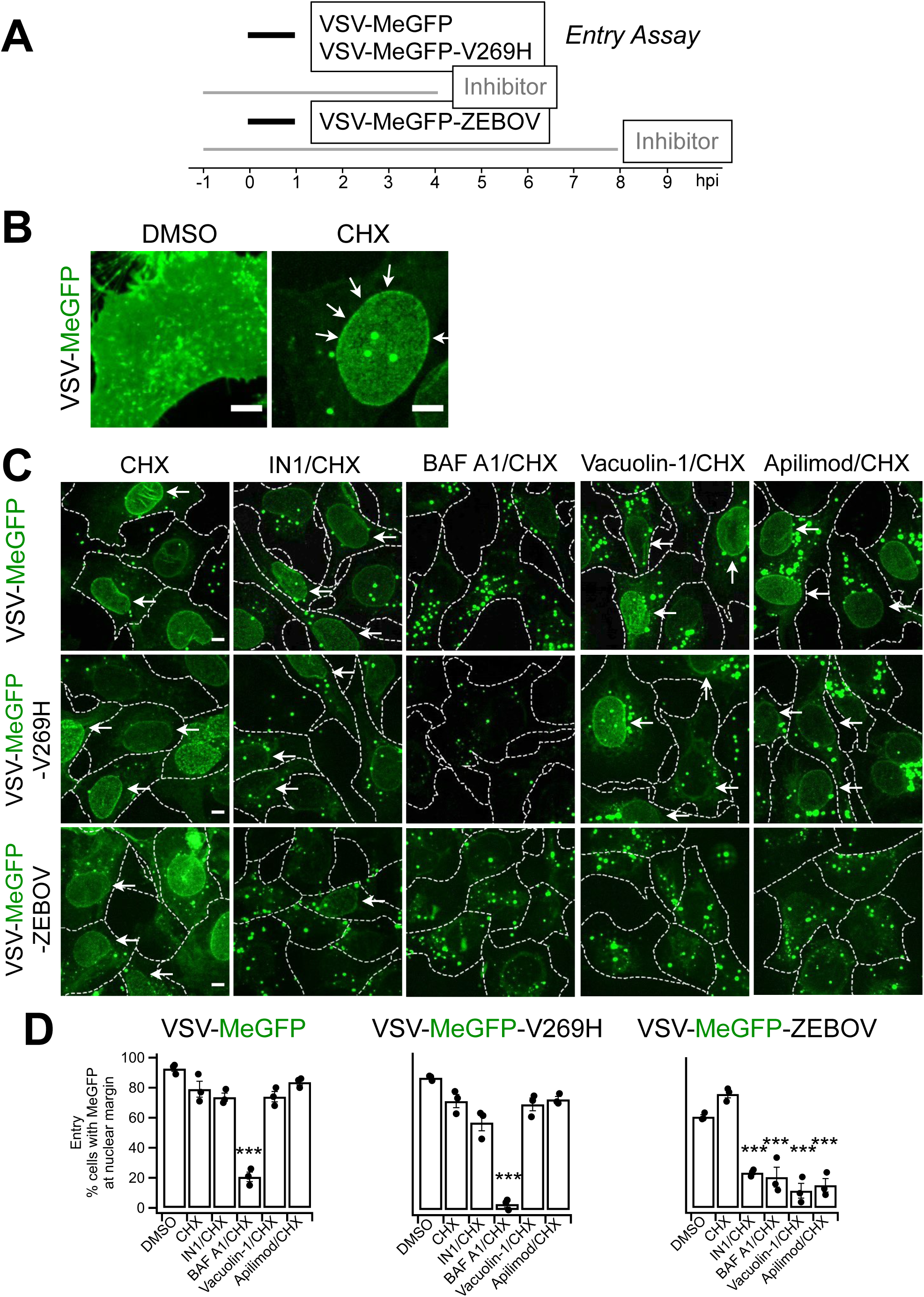
Apilimod and Vacuolin-1 inhibit VSV-MeGFP-ZEBOV. **(A)** Schematic of entry assay where SVG-A cells were infected with VSV-MeGFP (MOI = 4), VSV-MeGFP-V269H (MOI = 4), or VSV-MeGFP-ZEBOV (MOI = 4). Experiments were performed in the presence of 5 µg/mL cycloheximide (CHX) to prevent protein synthesis. Entry assay was based on the appearance of MeGFP fluorescence on the nuclear margin on a per cell basis, of fixed infected cells visualized by fluorescence microscopy. Staining the fixed cells with Alexa647 labeled wheat germ agglutinin identified the plasma membrane of each cell (dashed outlines in **C**). **(B)** Virus infection in the absence of CHX (left panel) resulted in the appearance of MeGFP fluorescence throughout the cell volume. The presence of CHX resulted in virus entry being observed by MeGFP fluorescence at the nuclear margin, which was released from incoming viral particles (right panel, white arrows). Scale bar indicates 10 µm. **(C)** Representative examples of maximum-Z projections images from the whole cell volume obtained with optical sections separated by 0.3 µm using spinning disc confocal microscopy. Scale bar indicates 10 µm. **(D)** Quantification of the number of cells with nuclear margin labeling from three independent experiments each determined from fields containing 59-90 cells (error bars show SEM). The statistical significance of the entry data was analyzed for statistical significance by one-way ANOVA and Tukey *post-hoc* test (***, P ≤ 0.001).

### Intracellular trafficking of virus particles in the presence of Apilimod or Vacuolin-1

Internalized virus particles traffic along the endocytic pathway to reach the endosomal compartment(s) from which membrane fusion and genome entry into the cytosol occur. To establish the identity of the endosomal compartments, we used genome-editing in SVG-A cells (**Figs. 3C, G and 4B, D**) to replace expression of a subset of proteins enriched in different endosomal compartments (the small GTPases Rab5c and Rab7a, EEA1, or NPC1) with their corresponding fluorescent chimeras obtained by fusion with TagRFP, mScarlet, or Halo (**Figs. 3B, E, F, I**, and **4C, E**). The lack of fluorescently tagged filipin (a cholesterol binder) in the endolysosomal compartment in the absence but not in the presence of U18666A, a potent inhibitor of NPC1 (**Fig 4F**), showed that NPC1-Halo remained active as a cholesterol transporter.

**Figure 3.**
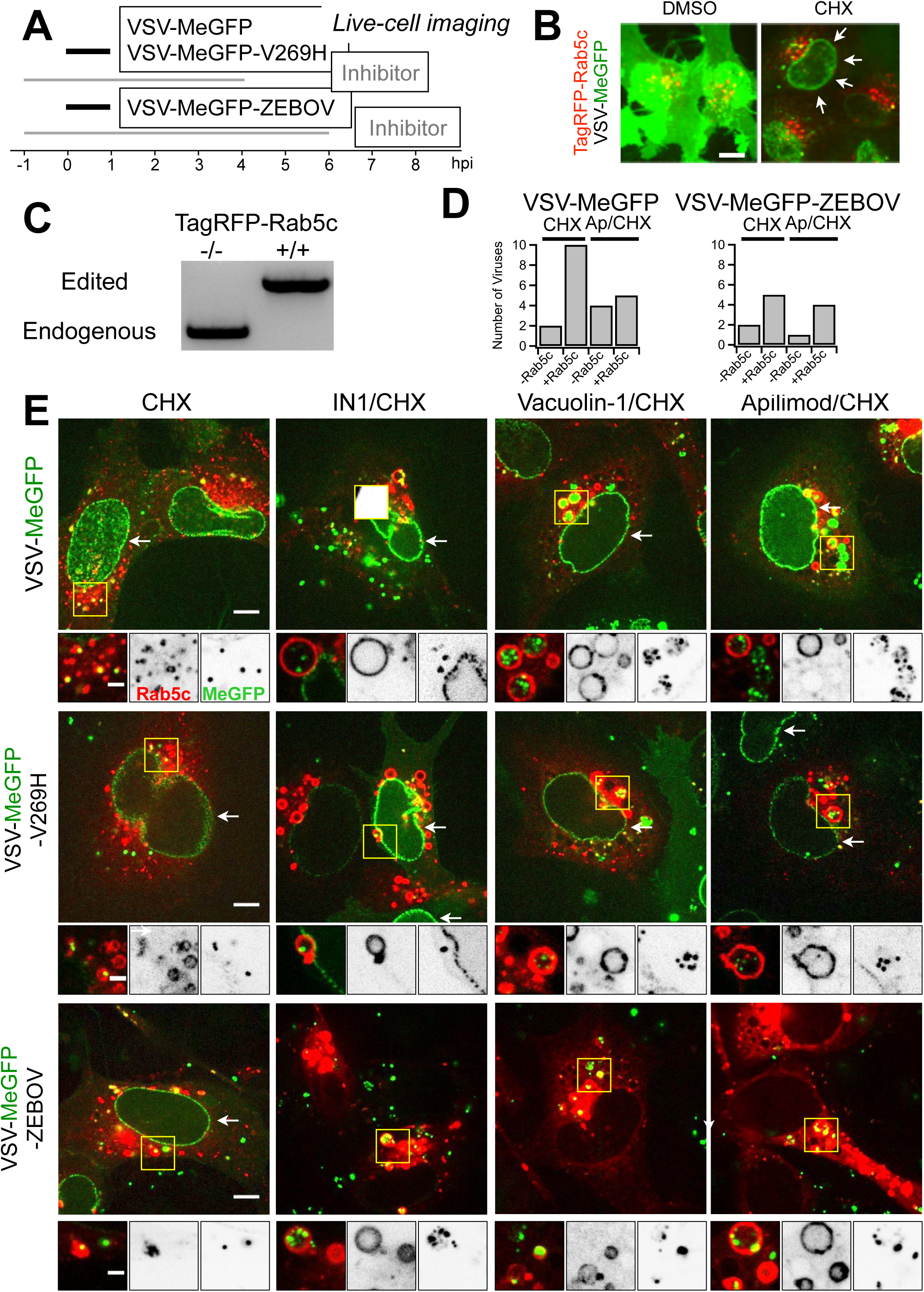
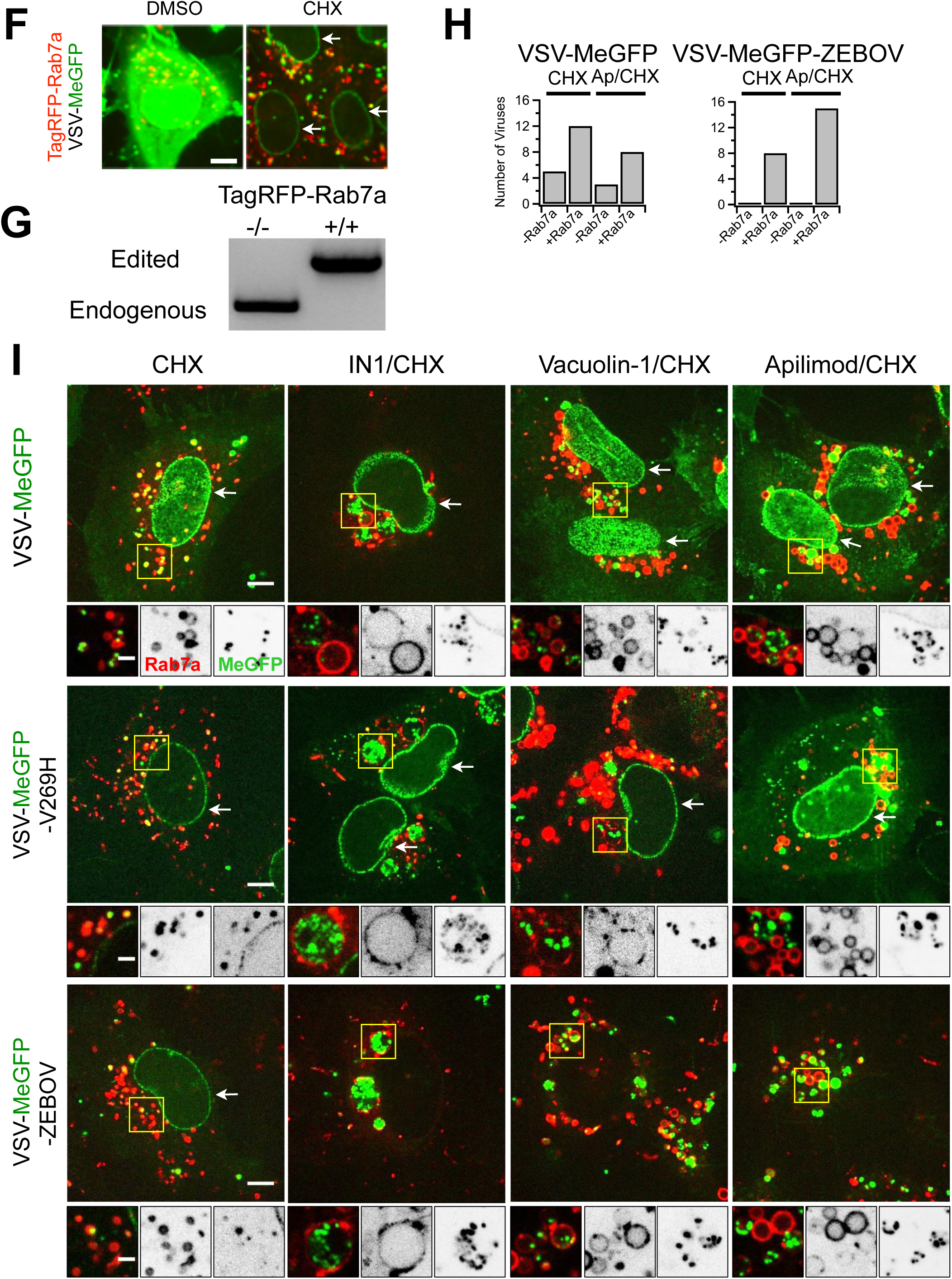
Endolysosomal traffic of VSV-MeGFP-ZEBOV in cells expressing TagRFP-Rab5c or TagRFP-Rab7a in the presence of Apilimod or Vacuolin-1. (Associated Videos 1 and 2). **(A)** Schematic of live cell imaging experiment using SVG-A cells expressing fluorescently tagged TagRFP-Rab5c or TagRFP-Rab7a. Cells were infected with VSV-MeGFP, VSV-MeGFP-V269H or VSV-MeGFP-ZEBOV (MOI = 4). Viruses trafficking (monitored with MeGFP) to the endo-lysosomal system (recognized by their labeling with TagRFP-Rab5c or TagRFP-Rab7a) and virus entry (established by MeGFP at the nuclear margin) were ascertained by live-cell florescence imaging using a spinning disc confocal microscope. **(B)** Visualization of VSV-MeGFP infection in TagRFP-Rab5c cells in the absence (left panel) or presence of CHX (right panel, white arrows) using live-cell imaging. Scale bar represents 10 µm. **(C)** Genomic PCR analysis of SVG-A cells showing biallelic integration of TagRFP into the *RAB5C* genomic locus by cotransfection of a plasmid coding for Cas9, a linear PCR product coding for the specific gRNAs targeting a region near the ATG codon of Rab5c under the control of the U6 promoter, and a template plasmid containing the RFP sequence flanked by 800 base pairs upstream and downstream of the targeted region (see materials and methods for more details) to generate a clonal gene-edited cell-line expressing TagRFP-Rab5c. **(D)** Quantification of VSV-MeGFP and VSV-MeGFP-ZEBOV colocalization with Rab5c containing endosomes in the presence of CHX together with absence or presence of 5 µM Apilimod depicted in (E). Data shows number of viruses that colocalized with endosomes containing or not Rab5c within the complete volume of the single cells depicted in (e). **(E)** Representative examples of maximum-Z projection images from four optical sections spaced 0.35 µm apart of virus entry without or with IN1, Vacuolin, or Apilimod for VSV-MeGFP (top), VSV-Me-GFP-V269H (middle), and VSV-MeGFP-ZEBOV (bottom). Each condition is in the presence of CHX. All viruses reach Rab5c-containing endosomes but only VSV-MeGFP-ZEBOV fails to penetrate in the presence of IN1, Vacuolin-1, or Apilimod. Scale bars are 10 µm. Insets correspond to a single optical section with scale bars of 3 µm. **(F)** Visualization of VSV infection in TagRFP-Rab7a cells in the absence of CHX (left panel) and entry in the presence of CHX (right panel, white arrows) with scale bar indicating 10 µm. **(G)** Genomic PCR analysis showing biallelic integration of TagRFP into the *RAB7A* genomic locus to generate a clonal gene-edited cell-line expressing TagRFP-Rab7a, using the same approach as used for *RAB5C*. **(H)** Quantification of VSV-MeGFP and VSV-MeGFP-ZEBOV colocalization with Rab7a containing endosomes in the presence of CHX with or without 5 µM Apilimod within the complete cell volumes in the images depicted in (I). **(I)** Representative examples of maximum-Z projection images from four optical sections spaced 0.35 µm apart of virus entry without or with IN1, Vacuolin, or Apilimod for VSV-MeGFP (top), VSV-Me-GFP-V269H (middle), and VSV-MeGFP-ZEBOV (bottom). All viruses reach Rab7a-containing endosomes but only VSV-MeGFP-ZEBOV fails to penetrate in the presence of IN1, Vacuolin-1 or Apilimod. Scale bars are 10 µm. Insets correspond to a single optical section with scale bars of 3 µm.

**Figure 4.**
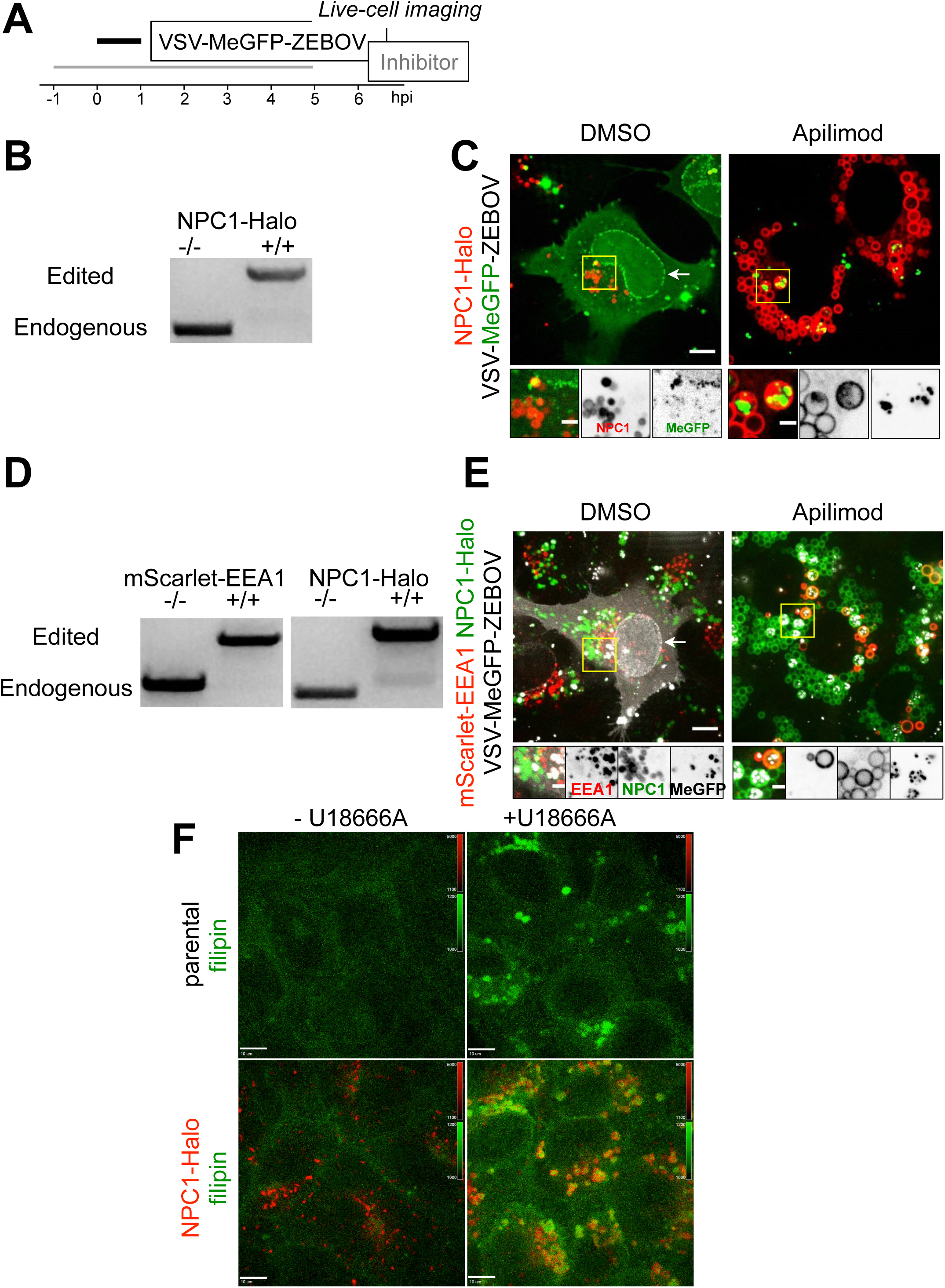
Endolysosomal traffic of VSV-MeGFP-ZEBOV in cells expressing NPC1-Halo or coexpressing mScarlet-EEA1 and NPC1-Halo in the presence of Apilimod. (Associated Video 3). **(A)** Schematic of live cell imaging experiment with gene-edited SVG-A cells expressing NPC1-Halo or NPC1-Halo together with mScarlet-EEA1. Halo was labeled with either JF549 or JF647. Cells were infected with VSV-MeGFP-ZEBOV (MOI = 3). **(B)** Genomic PCR analysis showing biallelic integration of Halo into the *NPC1* genomic locus to generate a clonal gene-edited cell-line expressing NPC1-Halo, using the same approach as for *RAB5C* and *RAB7A*. **(C)** Representative examples of maximum-Z projection images from four optical sections spaced 0.25 µm apart in the absence (left) and presence of Apilimod (right) showing that VSV-MeGFP-ZEBOV reached NPC1-Halo-containing endosomes even in the presence of Apilimod, while failing to penetrate and infect. Scale bar indicates 10 µm. Insets correspond to a single optical section with the scale bar indicating 3 µm. **(D)** SVG-A cells with genomic NPC1-Halo were further gene edited to contain EEA1 tagged with mScarlet. Genomic PCR analysis shows biallelic integration into the *EEA1* locus of mScarlet-EEA1 (left) and into the *NPC1* locus of NPC1-Halo (right). **(E)** Representative examples of maximum-Z projection images in the absence (left) and presence of Apilimod (right) showing that VSV-MeGFP-ZEBOV reached endosomes containing mScarlet-EEA1 and endosomes containing both mScarlet-EEA1 and NPC1-Halo in the presence of Apilimod, while failing to penetrate and infect. Scale bar indicates 10 µm. Insets correspond to a single optical section with scale bar indicating 3 µm. **(F)** Representative images of parental (top) and gene-edited SVG-A cells expressing NPC1-Halo (bottom) incubated with filipin III (naturally fluorescent polyene antibiotic that binds to cholesterol) in the absence (left) and presence of U18666A (right, NPC1 inhibitor of cholesterol export) showing NPC1-Halo is a functional cholesterol transporter.

Using live-cell spinning disk confocal microscopy (**Fig. 3, 4**), we monitored the presence of virus particles in the fluorescently tagged endosomes by colocalization with the fluorescent spots from the virus-incorporated MeGFP. We monitored entry by carrying out the experiments in the presence of cycloheximide, thus ensuring that any MeGFP fluorescent signal at the nuclear margin originated only from MeGFP molecules carried by incoming viral particles (**Fig. 3B, F**). All cells were maintained at 37°C throughout all phases of the experiment to ensure normal and undisturbed intracellular trafficking. All control experiments performed in the absence of inhibitors showed arrival of VSV-MeGFP, VSV-MeGFP-V269H, or VSV-MeGFP-ZEBOV virus particles to early (Rab5c and EEA1) (**Fig. 3E, 4E**) or late endosomes and lysosomes (Rab7a or NPC1) (**Fig. 3I, 4C, E**). MeGFP released from all viruses appeared at the nuclear margin, showing effective RNP release. NPC1, the receptor for VSV-MeGFP-ZEBOV entry is required for fusion from endosomes (2). The successful VSV-MeGFP-ZEBOV infection observed in the absence of drug in cells expressing NPC1-Halo alone or in combination with mScarlet-EEA1 indicates that NPC1-Halo is capable of facilitating infection and that VSV-MeGFP-ZEBOV trafficked to NPC1-Halo-containing endosomes.

Apilimod and Vacuolin-1 treatment of the SVG-A cells led to enlargement and vacuolization of their endosomes and lysosomes tagged with fluorescent EEA1, Rab5c, Rab7a or NPC1 (**Fig. 3-5**), in agreement with earlier PIKfyve ablation studies (13, 21). VSV-MeGFP and VSV-MeGFP-V269H (fluorescent dots, white) reached all tagged species of enlarged endolysosomes and successfully penetrated into the cytosol, as indicated by MeGFP at the nuclear margin (**Fig. 3E, I**). VSV-MeGFP-ZEBOV also trafficked to all tagged species of enlarged endolysosomes (**Fig. 3E, I**), often reaching one of the numerous NPC1-containing vacuoles enriched in EEA1 (**Figs. 4E** and **5B,C**). VSV-MeGFP-ZEBOV in EEA1-containing endosomes increased in the presence of Apilimod, as also reported for VLP ZEBOV (27). While able to reach NPC1-containing functional endosomes in cells treated with Apilimod (**Fig. 4C, E** and **5B, C**), VSV-MeGFP-ZEBOV failed to penetrate into the cytoplasm, as reflected by absence of MeGFP in the nuclear margin (**Fig. 2C, 3E, I, 4C,E and 5B**).

**Figure 5.**
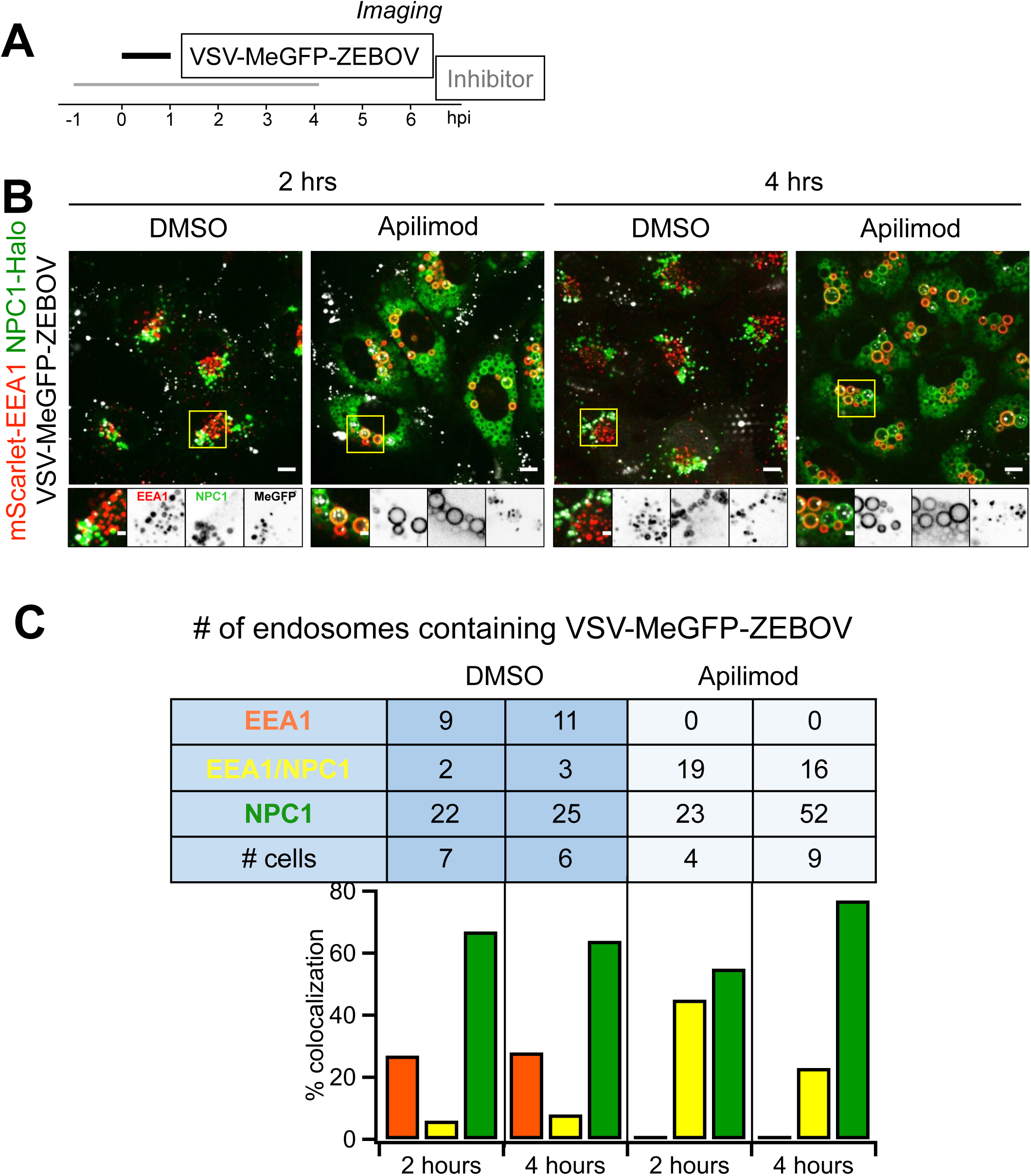
Extent of VSV-MeGFP-ZEBOV traffic into endosomes enriched in in NPC1-Halo or NPC1-Halo and mScarlet-EEA1. **(A)** Schematic of imaging experiment of VSV-MeGFP-ZEBOV trafficking in NPC1-Halo or NPC1-Halo and mScarlet-EEA1 gene edited SVG-A cells. **(B)** Representative examples of maximum-Z projection images from four optical sections spaced 0.25 µm apart in the absence and presence of Apilimod after 2 or 4 hours post infection. A large number of VSV-MeGFP-ZEBOV but not of VSV-MeGFP particles accumulated in the endosomes enlarged upon Apilimod treatment. **(C)** Quantification of VSV-MeGFP-ZEBOV colocalization with mScarlet-EEA1 alone, both mScarlet-EEA1 and NPC1-Halo, or NPC1-Halo alone 2 and 4 hours post infection, in the absence or presence of 5 µM Apilimod. Data obtained from complete cell volumes are presented as numbers and corresponding % colocalization<u>s</u> of VSV-MeGFP-ZEBOV particles associated with a given type of endosome.

### Apilimod blocks infection of VSV SARS-CoV-2

Using a recombinant vesicular stomatitis virus (VSV) expressing soluble eGFP (VSV-eGFP) where the glycoprotein (GP) was replaced with that of ZEBOV GP (VSV-eGFP-ZEBOV) or SARS-CoV-2 S (VSV-eGFP-SARS-Cov2), we inoculated MA104 cells with these chimera viruses and tested the effects of Apilimod on infection by flow cytometry (**Fig. 6A**). We found potent inhibition of VSV-eGFP-SARS-CoV-2 infection by Apilimod and confirmed that the compound also inhibits VSV-eGFP-ZEBOV infection (**Fig. 6B**). The dose-response curves indicated similar effects for VSV-eGFP-ZEBOV and VSV-eGFP-SARS-CoV-2 (IC50s ∼ 50 nM), in contrast to the absence of any detectable inhibition of VSV-eGFP infection, used here as a negative control.

**Figure 6.**
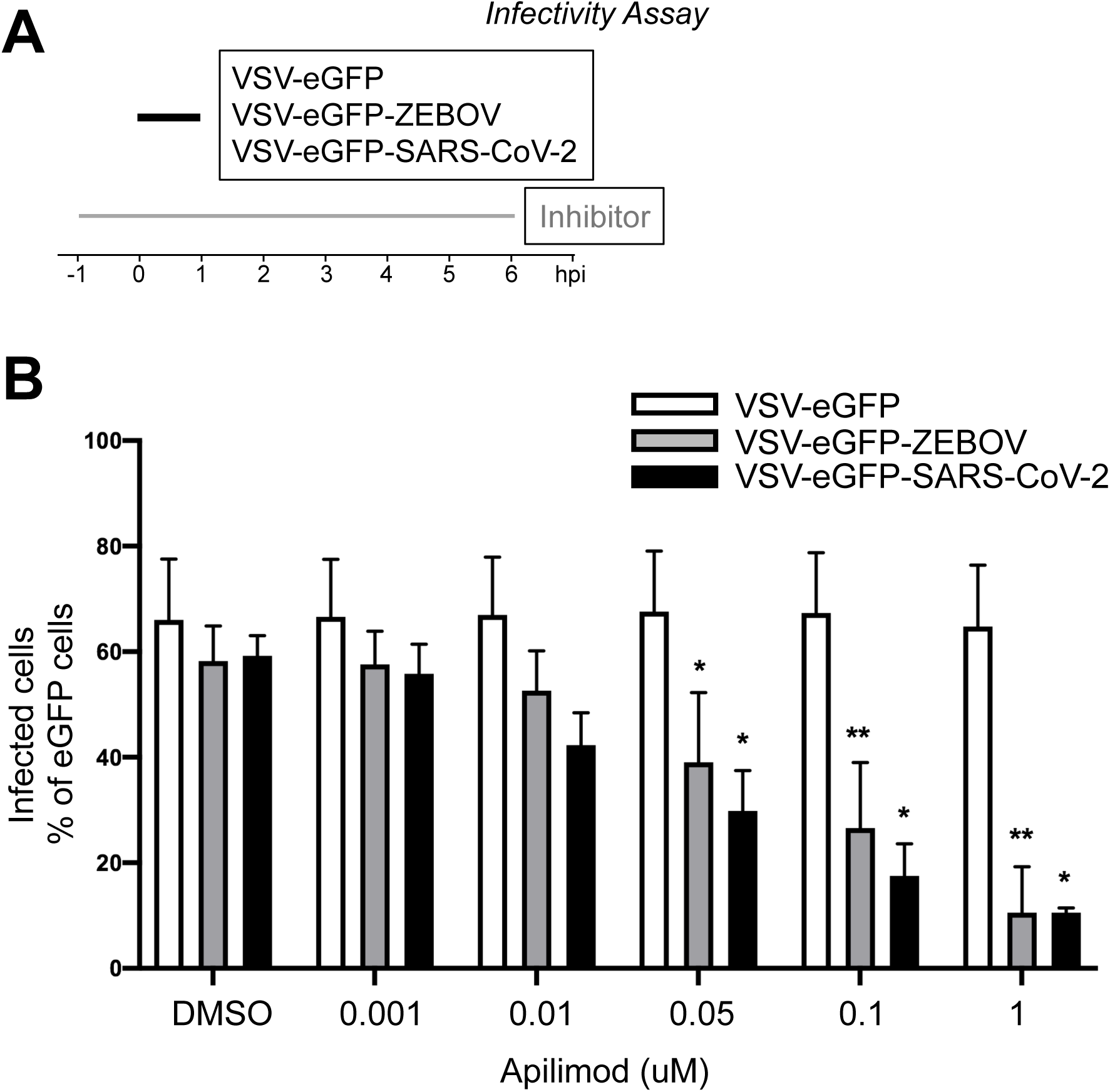
Apilimod and Vacuolin-1 inhibit infection of VSV-eGFP-SARS-CoV-2. **(A)** Schematic of infectivity assay of VSV-eGFP, VSV-eGFP-ZEBOV, and VSV-eGFP-SARS-CoV-2 in MA104 cells. MA104 cells were pretreated for 1 h with the indicated concentration Apilimod. Pretreated cells were inoculated with the indicated virus (MOI = 1) for 1 h at 37°C. At 6 hours post infection cells were harvested and the fraction of cell expressing eGFP cells quantified by flow cytometry. **(B)** Quantification of the infectivity is shown with averages +/- SEM from three independent experiments. Statistical significance was determined using a T-test (*, P ≤ 0.05; **, P ≤ 0.01).

### Apilimod blocks infection of SARS-CoV-2 virus

To test the effect of Apilimod on bona fide SARS-CoV-2 infection, we exposed Vero E6 cells to fully infectious SARS-CoV-2 (strain 2019-nCoV/USA-WA1/2020); after 24 h incubation, supernatants were harvested and tittered by focus-forming assay on a separate set of Vero E6 cells (**Fig. 7A**). Apilimod strongly inhibited SARS-CoV-2 infection, and the dose-response curve was similar or more potent than those observed for VSV-eGFP-ZEBOV or VSV-eGFP-SARS-CoV-2 (IC50s ∼ 10 nM) (**Fig. 7B**).

**Figure 7.**
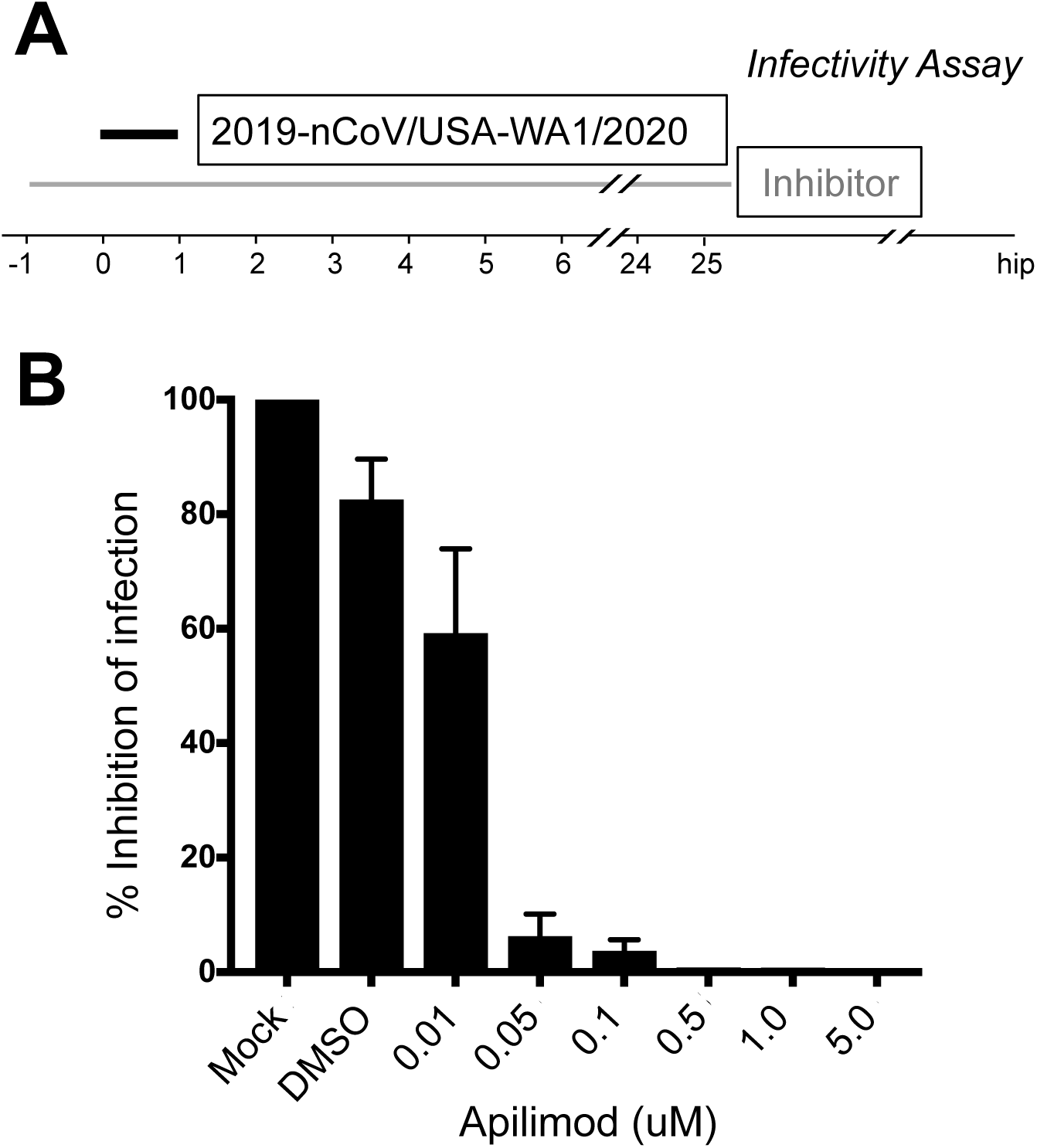
Apilimod inhibits infection of SARS-CoV-2 virus. **(A)** Schematic of infectivity assay of fully infectious Sars-CoV-2 (strain 2019-nCoV/USA-WA1/2020). Vero E6 cell monolayers were pretreated with medium containing DMSO or serial dilutions of Apilimod at the indicated concentrations. SARS-CoV-2 was diluted (MOI = 0.01) in Apilimod-containing medium and added to Vero E6 cells for 1 h at 37°C. After adsorption, the viral inocula were removed, and medium containing the respective concentration of Apilimod was reapplied. After 24 h incubation, supernatants were harvested and titrated by focus-forming assay on a separate set of Vero E6 cells. **(B)** Quantification of the infectivity is shown with averages +/- SEM from three independent experiments per condition and expressed as the percent infection relative to mock-treated SARS-CoV-2 infected cells.

## DISCUSSION

Coronaviruses, filoviruses, and arenaviruses have different replication strategies and unrelated surface glycoproteins that engage different receptor molecules during entry (1, 2, 5-8). Coronavirus and filovirus surface glycoproteins share a requirement for entry-associated proteolytic processing for activation as fusogens (1) Filoviruses require passage through low pH compartments where cathepsins are active. Coronaviruses may enter directly by fusion at the plasma membrane or following receptor mediated endocytosis. Cell entry of SARS-CoV and SARS-CoV-2 depends on the protease TMPRSS2 in conjunction with ACE2 (34-37), and when TMPRSS2 is present, the entry pathway becomes insensitive to cathepsin inhibition (34, 37, 38).

The common inhibition of viruses from all three groups by Apilimod is a consequence of perturbing their shared entry pathway. Moreover, it is not the cathepsin activity itself that these compounds affect, judging from the outcome of the assays with Apilimod and Vacuolin-1 showing they inhibit VSV chimeras bearing the surface glycoproteins of ZEBOV and LCMV and to a lesser extent LASV. Apilimod also inhibits infection of cells by VSV-SARS-CoV-2 as well as by authentic SARS-CoV-2; neither compound blocks infection by wild-type VSV. For VSV-ZEBOV, we have shown that the virus reaches a compartment enriched in NPC1, the ZEBOV co-receptor, and often also enriched in EEA1, but that it nonetheless fails to release internal proteins into the cytosol. Apilimod does not inhibit cathepsin (26) but Apilimod (39) and Vacuolin-1 (17, 23) can interfere with cathepsin maturation as evidenced by an increase in pro-cathepsin in treated cells; they do not influence endosomal pH (18, 26, 40) although other studies report Apilimod decreases cathepsin activity (41) and Vacuolin-1 increases pH (17, 23). Irrespective of this discrepancy, both Apilimod and Vacuolin-1 inhibit PI-3P-5-kinase (PIKfyve) (17, 19), a three-subunit complex (14) with a PI-3P-binding FYVE domain (10, 11) that recognizes the endosomal marker, PI-3-P. Functional ablation of this enzyme by genetic means (12, 15) gives rise to the same cellular phenotype as treatment with either compound (17-19). The similar dose-response curves for Apilimod inhibition of the ZEBOV and SARS-CoV-2 chimeras (IC50 of ∼ 50 nM) and of authentic SARS-CoV-2 virus (IC50 ∼ 10 nM) are in good agreement with the IC50 of ∼ 15 nM for Apilimod inhibition of PIKfyve *in vitro* (19). Thus, perturbing normal endosomal trafficking by inhibiting PIKfyve activity suggests it is the mechanism by which Apilimod and Vacuolin-1 block entry of such a diverse set of viral pathogens.

One of the most striking consequence of PIKfyve inhibition, and hence of PI-3,5-P_2_ restriction in endosomal membranes, is the swelling of endosomes into small, spherical vacuoles - the phenomenon that gave Vacuolin-1 its name (18). Our imaging data with VSV-MeGFP-ZEBOV chimeras show that the virus particles accumulating in these structures, many of which also contain the NPC1 co-receptor (2, 42), often appear to be relatively immobile and adjacent to the endosomal limiting membrane. One possible explanation is that when a virion reaches these distended endosomes, it can bind or remain bound to the limiting membrane, but not fuse. Another is that virions may fuse with smaller intraluminal vesicles in the endosomal lumen (43), but that PI-3,5-P2 depletion prevents back fusion of these vesicles with the endosomal limiting membrane and inhibits release into the cytosol of the viral genome.

Inhibition of infection by authentic SARS-CoV-2 shows that the blocked release of the viral genome from a vacuolated endosome is independent of the shape, size, and distribution of spike protein on the virion. The assay we used to determine effects on infectivity of authentic virus measured release of virions after multiple rounds of infection, rather than entry, which we monitored in the VSV-SARS-CoV-2 experiments by detecting eGFP synthesis in the cytosol. Nevertheless, the IC50 of Apilimod in experiments with authentic virus is remarkably similar (or even more potent) to that obtained with chimeric VSV-SARS-CoV-2.

Although cathepsin L inhibitors block SARS-CoV and SARS-CoV-2 infection in cell culture (4, 5), they have less pronounced effects when tested in animals (44). This may because another protease, TMPRSS2 on the surface of cells in relevant tissues, appears to prime SARS-CoV (44) and SARS-CoV-2 (37) spike proteins for efficient entry. As the effectiveness of Apilimod and Vacuolin-1 does not depend on cathepsin inhibition, their capacity to block entry of several distinct families of viruses is likely to be independent and downstream of the protease that primes their surface glycoprotein for fusion. Phase I and phase II clinical trials have shown that Apilimod is safe and well-tolerated (24, 25, 29, 30). The trials were discontinued because of lack of effectiveness against the autoimmune condition for which the drug was tested. We suggest that one of these compounds, or a potential derivative, could be a candidate broad-spectrum therapeutic for several emerging human viral pathogens, including SARS-CoV-2.

---

**Video 1. Apilimod doesn’t inhibit VSV-MeGFP entry**. Maximal Z-projection from four optical sections separated 0.25 µm apart of SVG-A cells gene-edited to express TagRFP-Rab5c imaged by spinning disc confocal microscopy every 3 seconds for 3 min. Cells were infected with VSV-MeGFP (MOI = 4) in the presence of CHX with or without 5 µM Apilimod and imaged ∼ 3-4 h post-infection.

**Video 2. Apilimod inhibits VSV-MeGFP-ZEBOV entry**. Maximal Z-projection from four optical sections separated 0.25 µM apart of SVG-A cells gene-edited to express TagRFP-Rab5c imaged by spinning disc confocal microscopy every 3 seconds for 3 min. Cells were infected with VSV-MeGFP-ZEBOV (MOI = 3) in the presence of CHX with or without 5 µM Apilimod and imaged ∼ 6-7 h post infection.

**Video 3. Apilimod inhibits VSV-MeGFP-ZEBOV entry**. Maximal Z-projection from four optical sections separated 0.25 µM apart of SVG-A cells gene-edited to express NPC1-Halo imaged by spinning disc confocal microscopy every 3 seconds for 3 min. Cells were infected with VSV-MeGFP-ZEBOV (MOI = 3) with or without 5 µM Apilimod and imaged ∼ 5 h post infection.

## MATERIAL AND METHODS

### Cell culture

Human astroglial SVG-A derived cells (a kind gift from Walter J. Atwood) were grown at 37°C and 5% CO_2_ in Minimum Essential Medium (MEM) (10-010-CV; Corning) supplemented with 10% heat inactivated fetal bovine serum (S11150; Atlanta Biologicals), penicillin and streptomycin (1406-05-9; VWR International). African Green Monkey kidney epithelial MA104 cells (a kind gift from Siyuan Ding, WUSM) were grown at 37°C and 5% CO_2_ in Medium 199 supplemented with 10% heat inactivated fetal bovine serum. Vero C1008 [Vero 76, clone E6, Vero E6] (ATCC CRL-1586) cells were cultured in Dulbecco’s Modified Eagle Medium (DMEM) supplemented with 10% fetal bovine serum, and penicillin and streptomycin. Vero CCL-81 (ATCC CCL-81) cells were maintained in DMEM supplemented with 10% FBS, 10mM HEPES pH 7.4, 1% Glutamax, and penicillin/streptomycin.

### Reagents

Vacuolin-1 (18) was custom synthesized; Apilimod (HY-14644) was from MedChem Express, IN1 was a kind gift from Dr. N. Gray (33), U-18666A (10009085), and Filipin III (70440) were from Cayman Chemical, Bafilomycin A1 (B1793-2UG) was from Sigma-Aldrich, Cycloheximide (239764) was from Calbiochem, and wheat germ agglutinin conjugated with Alexa Fluor®-647 (W32466) was from ThermoFisher.

### Viruses

Recombinant VSV (Indiana serotype) expressing MeGFP alone which initiates fusion at pH<6.2 (VSV-MeGFP) (45) (or in combination with V269H GP, VSV-MeGFP-V269H), RABV GP (VSV-MeGFP-RABV) (46), LASV GP (VSV-MeGFP-LASV) (7), LCMV GP (VSV-MeGFP-LCMV), Zaire EBOV GP (VSV-MeGFP-ZEBOV) (47) or SARS-CoV-2 S Wuhan-Hu-1 strain (VSV-eGFP-SARS-CoV-2 – description to be published elsewhere) were used for infection, entry and live cell imaging assays. All viruses were generated and recovered according to (48).

SARS-CoV-2 strain 2019-nCoV/USA-WA1/2020 was obtained from the Centers for Disease Control and Prevention (gift of Natalie Thornburg). Virus was passaged once in Vero CCL81 cells (ATCC) and titrated by focus-forming assay also on Vero E6 cells.

### Genome-editing

Individual cell lines of SVG-A were gene edited in both alleles using the CRISPR/Cas9 system to incorporate fluorescent tags into the N-terminus of Rab5c (TagRFP), Rab7a (TagRFP), EEA1 (mScarlet) or the C-terminus of NPC1 (Halo). The NPC1-Halo expressing cells were further gene edited to incorporate mScarlet-EEA1 creating SVG-A cells simultaneously expressing mScarlet-EEA1 and NPC1-Halo.

A free PCR strategy (49, 50) was used to generate small guide RNAs (sgRNA) with target sequences for either Rab5c, Rab7a, NPC1, or EEA1.

The genomic DNA fragment of Rab5c, Rab7a, NPC1, or EEA1 genes fused with either TagRFP, Halo, or mScarlet were cloned into the pUC19 vector (donor constructs) which then served as homologous recombination repair templates for the Cas9 enzyme-cleaved genomic DNA. Donor constructs were obtained by a ligation of PCR amplification products from the genomic DNA fragments, TagRFP, Halo, and mScarlet sequences. Primers F1-R1 and F3-R3 amplified approximately 800 base pairs of genomic sequences upstream and downstream of the start codon of Rab5c, Rab7a or EEA1, or the stop codon of NPC1, respectively. Primers F1 and R3 contain sequences complementary to the pUC19 vector linearized using the SmaI restriction enzyme (lower case in the primer sequences). The TagRFP sequence containing the GGS peptide linker was amplified using primers F2-R2 from a TagRFP mammalian expression plasmid used as a template. The F2 primer contains complementary sequences to the 3’ end of the F1-R1 fragment, while the F3 primer contains complementary sequences to the 3’ end of the TagRFP sequences. Primer sequences used to generate the sgRNAs and corresponding genomic fragments are listed in Table I; primers used for screening are listed in Table II.

**Table I.**
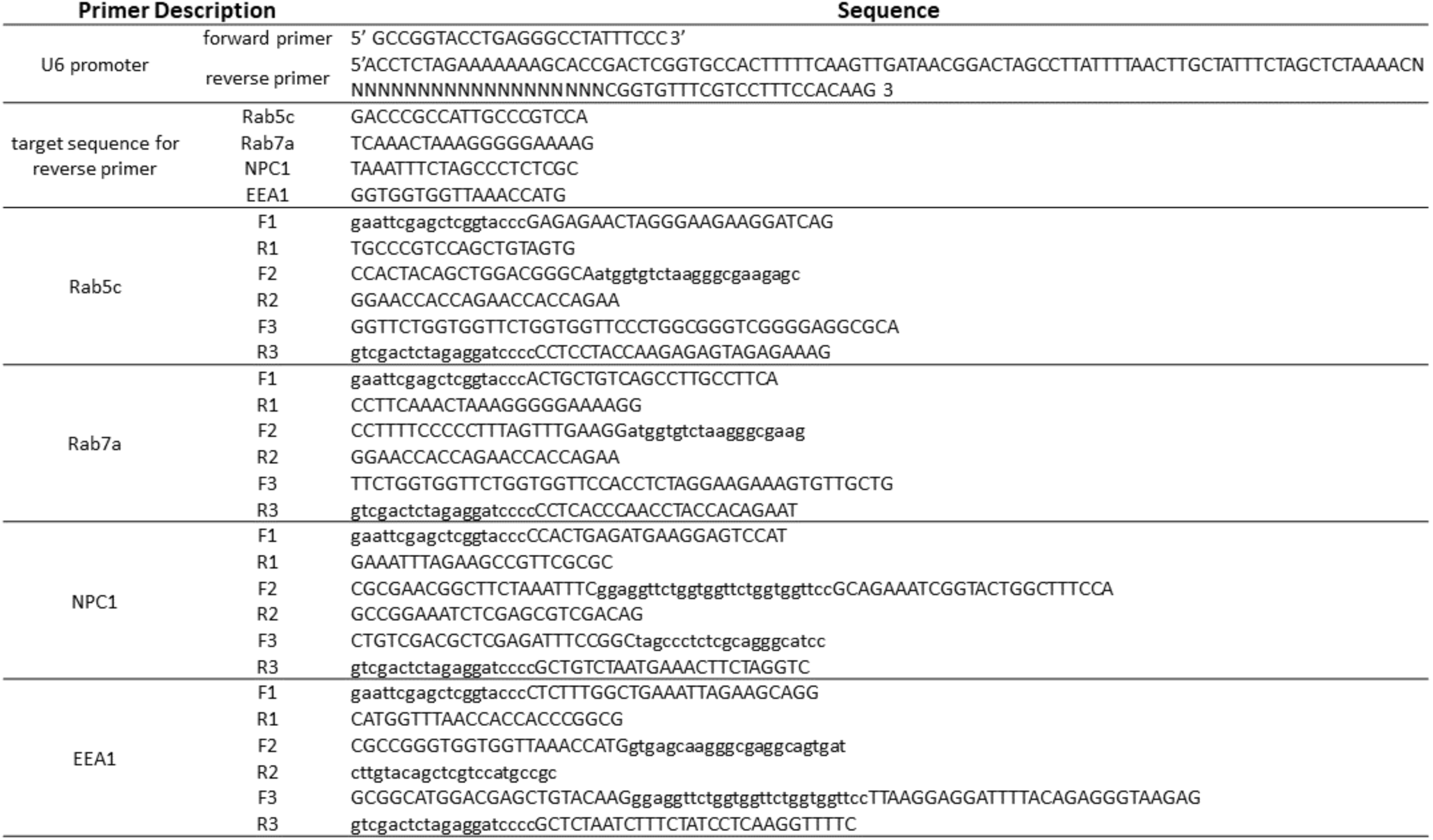
Primer sequences used to generate the sgRNAs and corresponding genomic fragments.

**Table II.**
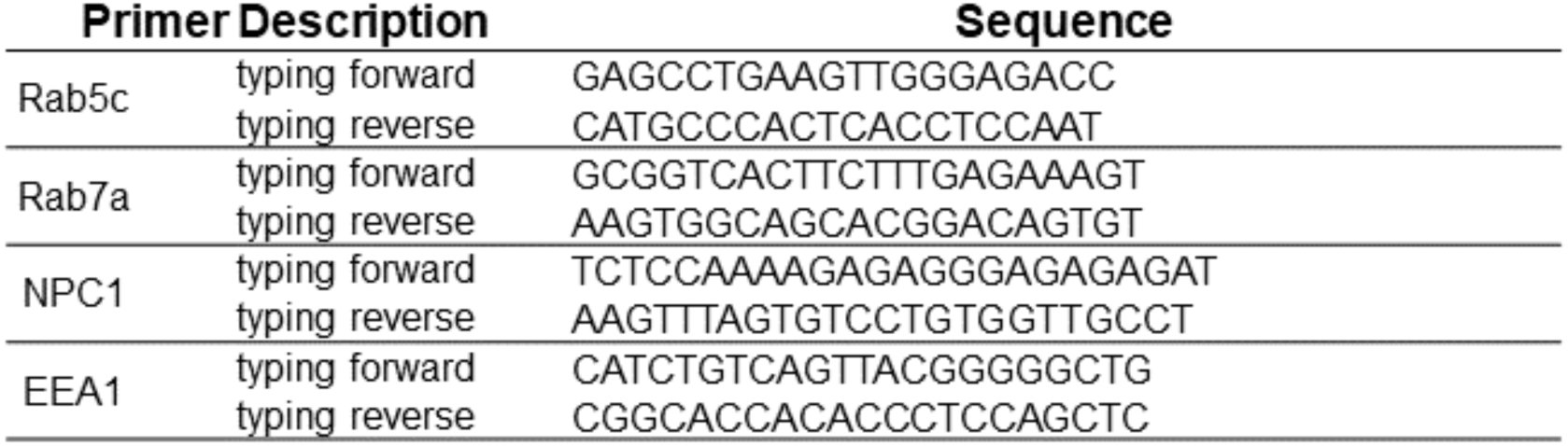
Primer sequences used for screening.

PCR products (fragments F1-R1, F2-R2, and F3-R3) were subjected to electrophoresis in 1% agarose and gel purified using a purification kit from Zymogen. The PCR fragments were cloned into the linearized pUC19 vector using the Gibson Assembly Cloning Kit (E5510S; New England Biolabs).

SVG-A cells (1.5 × 10^5^ cells) were co-transfected with 0.8 µg of *Streptococcus pyogenes* Cas9, 0.8 µg free PCR product coding for the target sgRNA, and 0.8 µg pUC19 vector using Lipofectamine 2000 reagent (Invitrogen) according to the manufacturer’s instructions. Transfected cells were grown for 7 to 10 days and sorted for TagRFP, Halo, or mScarlet expression using fluorescence-activated cell sorting (FACS) (SH-800S; Sony). Prior to FACS, NPC1-Halo cells were labeled for 15 minutes with Janelia Fluor^™^ 647 (JF647). Single cells expressing the desired chimera were isolated, clonally expanded, and then screened by genomic PCR for TagRFP, Halo, or mScarlet insertion into both alleles.

### Infection assays

SVG-A cells were plated at about 30-40% confluency into 24-well plates and incubated for 1 day at 37°C and 5% CO_2_. At the start of the experiment, cells were incubated with the indicated drug or DMSO at 37°C for one hour. Following this, cells were incubated for 1 h at 37°C with VSV, VSV-MeGFP-V269H, VSV-MeGFP-RABV, VSV-MeGFP-LASV, VSV-MeGFP-LCMV or VSV-MeGFP-ZEBOV in drug or DMSO-containing infection medium (α-MEM, 50mM HEPES, 2% FBS). Cells were then washed to remove non-adsorbed viruses and further incubated at 37°C in medium containing the drug or DMSO with experiments ending at the indicated times by fixation with 3.7% formaldehyde in PBS. Fluorescent intensity from 20,000 single cells from a single round of infection was determined by flow cytometry using a BD FACSCanto^™^ II equipped with DIVA software package.

MA104 cells were pretreated for 1 h with the indicated concentration Apilimod or DMSO. Pretreated cells were inoculated with VSV-eGFP, VSV-eGFP-ZEBOV or VSV-eGFP-SARS-CoV-2 at an MOI = 1 (based on titers in MA104 cells) in the presence of Apilimod or DMSO for 1 h at 37°C. Six to 8 h post infection, cells were collected and fixed in 2% PFA and then subjected to flow cytometry. The percentage of GFP cells was determined using FlowJo software (Tree Star Industries, Ashland, OR).

Vero E6 cell monolayers were pretreated for 1 h at 37°C with serial dilutions of Apilimod at the indicated concentrations. Next, SARS-CoV-2 was diluted to an MOI of 0.01 focus-forming units (FFU)/cell in Apilimod-containing medium and added to Vero E6 cells for 1 h at 37°C. After adsorption, cells were washed once with PBS, and medium containing the respective concentration of Apilimod was added. Cell were incubated for a 24 h at 37°C, and at which time cell culture supernatants were removed and used for determination of viral titer by focus forming assay.

### SARS-CoV-2 focus forming assay

Cell culture supernatants from virus-infected cells were diluted serially 10-fold and added to Vero E6 cell monolayers in 96-well plates and incubated at 37°C for 1 h.

Subsequently, cells were overlaid with 1 % (w/v) methylcellulose in MEM supplemented with 2% FBS. Plates were harvested 30 h later by removing overlays and fixed with 4% paraformaldehdye in PBS for 20 min at room temperature. Plates were washed and sequentially incubated with 1 µg/mL of CR3022 anti-spike antibody (51) and HRP-conjugated goat anti-human IgG in PBS supplemented with 0.1% saponin and 0.1% BSA. SARS-CoV-2-infected cell foci were visualized using TrueBlue peroxidase substrate (KPL) and quantitated on an ImmunoSpot microanalyzer (Cellular Technologies). Data were processed using Prism software (GraphPad Prism 8.0) and viral titers are reported as percent inhibition relative to mock-treated SARS-CoV-2 infected cells.

### Entry assay and intracellular traffic

SVG-A cells plated on glass #1.5 coverslips at about 30-40% confluency one day prior to experiment were treated with drug or DMSO for 1 h at 37°C. Following this, cells were incubated at 37°C with VSV, VSV-MeGFP-V269H, VSV-MeGFP-RABV, VSV-MeGFP-LASV, VSV-MeGFP-LCMV or VSV-MeGFP-ZEBOV in drug or DMSO containing infection medium. After this, cells were washed then further incubated in medium containing the drug or DMSO at 37°C with the experiment ending at the indicated time by fixation for 20 min at room temperature with 3.7% formaldehyde in PBS. This was followed with a 10-min incubation of 5 µg/mL of Alexa647-labeled wheat germ agglutinin in PBS to label the outline of the cells.

Cells were imaged using a spinning disk confocal microscope with optical planes spaced 0.3 µm apart (52). The entry assay scored the presence of MeGFP at the nuclear margin in each cell. Trafficking of viruses to endosomal compartments was observed using live-cell imaging using the spinning disc confocal microscope. Chemical fixation tends to eliminate the large endolysosomal vacuoles generated by Vacuolin-1 or Apilimod and reduces the colocalization with viral particles contained within. Time series with images taken every 3 seconds for 3 min in a single optical plane with the appropriate fluorescent channels (52) were acquired from non-fixed samples imaged at the end of the experimental period. For experiments containing NPC1-Halo, the Halo-tagged cells were labeled with either 250 nM JF549 or JF647 dye in media for 30 min at 37°C. Following labeling, cells were washed three times with media. The microscope was operated using the Slidebook 6.4 software package (3I) and images were displayed also using this software.

### Statistical tests

To compare the means from cells with different treatments, one-way ANOVA and the *post-hoc* Tukey test analysis were used to take into account unequal sample sizes as indicated in figure legends.

## ACKNOWLEDGEMENTS

We thank Walter J. Atwood for providing the parental SVG-A cells, Eric Marino, Justin H. Houser and Tegy John Vadakkan for maintaining the spinning disc confocal microscope (T.K. Lab), Marina Cella and Erica Lantelme from the flow cytometry facility, Department of Pathology and Immunology, WUSM for help with flow cytometry, Stephen C. Harrison for helpful discussions and editorial assistance and Alex J. B. Kreutzberger for editorial help. This research was supported by National Institutes of Health funding (AI109740) to S.P.W. and T.K, by MIRA NIH award (GM130386) to T.K., by National Institutes of Health funding (AI059371) and unrestricted funds from WUSM to S.P.W. and by (75N93019C00062 and R01 AI127828) to M.S.D.

## AUTHOR CONTRIBUTIONS (NAMES GIVEN AS INITIALS)

T.K., S.P.W., and M.S.D. were responsible for the overall design of the study; Y.K. carried out virus infection, entry and imaging experiments and prepared figures in the lab of T.K. (Fig. 1-5). P.W.R. designed, generated and characterized VSV-eGFP-SARS-CoV-2 and Z.L. carried out VSV-chimera infection experiments in the lab of S.P.W. (Fig. 6). J.B.C. and R.E.C. carried out the experiments with authentic SARS-CoV-2 under BSL3 conditions in the lab of M.S.D. (Fig. 7). Recombinant viruses were generated and characterized by D.K.C., S.P., M.R and T.S. in the lab of S.P.W; T.K. drafted the manuscript and editorially reviewed it in close association with SP.W. and M.S.D; the authors commented on the manuscript.

